# Hatching asynchrony and fitness in a neotropical seabird: second-hatched individuals from highly asynchronous broods are less fit but only during the rearing period

**DOI:** 10.1101/2023.05.11.540419

**Authors:** Santiago Ortega, Cristina Rodríguez, Hugh Drummond

## Abstract

Early-life conditions are important determinants of phenotype and fitness. In birds, hatching asynchrony can generate differences in early-life conditions within a brood, which, in turn, can have far reaching fitness consequences for offspring, particularly so for later-hatched nestlings. A plethora of literature has examined consequences of hatching asynchrony during the nestling phase; however, long-term effects remain poorly understood. Using a 33-year population study of the Blue-footed booby (*Sula nebouxii*) off the Pacific Coast of Mexico, we show that the level of hatching asynchrony affects early- life survival of second-hatched nestlings. Junior boobies from highly asynchronous broods died at younger ages during the rearing period and were less likely to fledge compared to first-hatched offspring. However, level of hatching asynchrony did not have long-term fitness effects on either senior- or junior juveniles or adults. Our results provide insight into how parentally imposed natal environments affect early-life survival and late-life fitness traits in a long-lived seabird.

**Highlights:**

- Parentally imposed natal conditions can have far reaching fitness consequences for offspring.
- In birds, hatching asynchrony can produce size hierarchies within a brood.
- The effects of natural variation in hatching asynchrony on fitness is poorly understood.
- High levels of hatching asynchrony are detrimental for junior booby nestlings.
- Hatching asynchrony does not affect fitness in the juvenile or adult periods.

## Introduction

In vertebrates, numerous experiments have shown that early-life conditions can alter an individual’s physiology, behaviour, morphology, and life history traits during adulthood (Grace et al., 2017; Haywood & Perrins, 1992; Kärkkäinen et al., 2021; Lemaître et al., 2015; Mumme et al., 2015; Nettle et al., 2017). In particular, natural early-life adversity has been linked to harmful consequences on fitness components and senescence rates in mammals (e.g., (Nussey et al., 2007; Pigeon et al., 2017; Plard et al., 2015), and in 5 of 9 bird species studied in nature, adults that experienced poor weather, habitats, prey availability, or parental care, or fledged late in the season showed reduced lifetime reproductive success (reviewed in (Drummond & Ancona, 2015)). Furthermore, experiments and a single descriptive field study have documented intergenerational effects of a poor start: nutritional and social stress in infancy can negatively affect the quality and viability of an animal’s eventual offspring (Burton & Metcalfe, 2014; Drummond & Rodríguez, 2013; Naguib & Gil, 2005). These findings reveal the important role that early- life conditions can play in determining later-life phenotype and reproductive performance.

In birds, differences in early-life conditions can even arise within a brood, particularly when parents begin incubation before clutch completion, causing asynchronous hatching that can span from hours to days (Clark & Wilson, 1981). Hatching asynchrony is recognized as an evolved –and female controlled– strategy (reviewed in (Magrath, 1990) that creates competitive inequality among nestlings, enables parents to reduce brood size to a number closely linked with food supplies (Lack, 1968), shortens the time offspring spend in the nest exposed to predators (Clark & Wilson, 1981), or spaces out the energy costs of rearing several offspring (Hussell, 1972).

Due to their relatively small size and delayed motor development, later-hatched nestlings tend to be competitively inferior to their older siblings and suffer short-term fitness penalties. For instance, last-hatched chicks of herring gull *Larus argentatus* broods with experimentally increased hatching asynchrony, showed delayed motor coordination and were less likely to survive the first 10 days of life –presumably due to their incapacity to escape predators– compared to chicks from unmanipulated broods (Hillström et al., 2000). Furthermore, later-hatched nestlings can grow more slowly and fledge in poorer body condition –a fitness prospect trait in birds, that is, a trait closely linked to late-life fitness (Morrison et al., 2009). For example, in the collared flycatcher, *Ficedula albicollis*, smaller nestlings –seemingly later-hatched ones– grew more slowly and had shorter wing feathers than nestmates from more synchronous broods (Szöllosi et al., 2007). Interestingly, later-hatched chicks can sometimes catch up –in growth and size– with their older siblings during the rearing period (e.g. (Clotfelter et al., 2000). However, catching-up can come at the expense of self-maintenance. For example, later-hatched great tit *Parus major* nestlings, despite an initial size handicap, grew at a similar rate and achieved a mass at fledging comparable to their older siblings, but exhibited greater telomere erosion (Stier et al., 2015), implying fitness penalties later in life (Metcalfe & Monaghan, 2001).

Staggered hatching is widespread among birds (Clark & Wilson, 1981) but, little is known about the effects of natural continuous variation in hatching asynchrony on the nestling, juvenile, and adult stages of either wild or captive birds, since most studies narrow down the level of asynchrony to two or three simple categories. For example, nestlings of wild pied flycatchers *Ficedula hypoleuca* from both experimentally synchronous and naturally asynchronous broods were found to differ in early survival and fitness prospect traits: synchronous broods grew faster, showed a higher body mass in the first weeks of life, and both nestmates showed similar telomere lengths at ages 5 and 12 days compared with naturally asynchronous broods (Kärkkäinen et al., 2021). Furthermore, a study of captive female zebra finches *Taeniopygia guttata* that were raised in asynchronous or experimentally synchronous broods, showed late-life effects of hatching asynchrony: elder females from asynchronous broods deposited more carotenoids and retinol in the egg yolk – a proxy of fecundity– than females from synchronous broods (Mainwaring et al., 2012). However, as long-term monitoring of broods with different spans of asynchrony is hard to achieve in the field, the effects of natural variation in hatching asynchrony on both early- survival, fitness prospects traits and late-life fitness have not, to our knowledge, been explored.

Here, we evaluated whether hatching asynchrony –across its natural span– has short- and long-term fitness consequences in the blue-footed booby *Sula nebouxii*, a long-lived neotropical seabird with extraordinarily long laying and hatching intervals. In our study population, females commonly lay two similar sized eggs (Drummond et al., 1986) that hatch at a median interval of 5 days (Ortega et al., unpublished data). The initial size disparities due to hatching asynchrony contribute to formation of a stable dominance relationship between nestmates during the first few weeks, in which the junior chick is overwhelmed by fierce and persistent pecking and learns to submit routinely to its senior sibling (Drummond et al. 1986, Drummond and Osorno 1992). Consequently, juniors suffer restricted food ingestion, slow growth and high circulating corticosterone during much of the nestling period (Drummond et al., 1991; Mora et al., 1996), and if food provision declines, aggressive brood reduction can occur: the dominant sibling ramps-up its aggression, killing its sibling or expelling it from the nest (Drummond & Garcia Chavelas, 1989).

Cross-fostering in the field showed that reduction or enhancement of hatching asynchrony can alter the agonism and the growth of broodmates. At 11-20 days, senior chicks from synchronous broods (mean hatching span of 0.16 days) pecked their younger nestmates 3 times more frequently than their counterparts in broods with average asynchrony (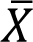 span of 4.08 days), and senior chicks from doubly asynchronous broods (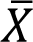 span of 7.53 days) were more aggressive than those from averagely asynchronous broods, pecking juniors five times as frequently when the latter were 0-10 days old. Interestingly, no negative effects of this elevated nestmate aggression on either seniors’ or juniors’ survival during their first month of life were detected, but juniors from doubly asynchronous broods were 11% lighter than juniors in averagely asynchronous broods at 10 days of age (Osorno & Drummond, 1995).

In this population of boobies, chicks do not become independent until after age 3 months and fitness consequences of hatching order and brood sexual composition beyond the nestling period appear to be few and minor. Junior recruits from broods of two produced low-viability fledglings (i.e., unlikely to recruit), albeit only in their first 3 years of life when few fledglings are produced (Drummond & Rodríguez, 2013). In addition, junior females that grew up with an elder brother that they outgrew (females being larger than males) showed a trivial (0.08%) deficit in annual hatching success, but no other negative impacts on their life-history trajectories or fitness (Drummond et al., 2022). Otherwise, pairs of senior- and junior fledglings of the Isla Isabel population appear to be equivalent as juveniles and adults: they do not differ at any age in defensive behaviour, survival or reproductive success (e.g., (Drummond et al., 2011; Drummond & Rodríguez, 2013; Sánchez-Macouzet & Drummond, 2011).

Hitherto, effects of natural variation in hatching asynchrony on fitness of blue-footed boobies have not been sought. Potentially, the span of hatching asynchrony between nestmates could cause differences in the performance of senior and junior individuals during the nestling, juvenile and adult periods. Since aggression by the elder sibling is considerably more intense in broods with either near-zero or doubled levels of hatching asynchrony, we predicted inverted-U relationships: (1) during the nestling period, fledging success, age at death and fledging weight should be lower for junior nestlings in broods with either low or high levels of hatching asynchrony than in broods with ∼ 5 days asynchrony. (2) After fledging, the probability and age of recruitment of junior juveniles from broods with low or high levels of hatching asynchrony should be lower because fledglings reared under stressful ecological conditions tend to recruit earlier (Ancona & Drummond, 2013). Finally, (3) we expected that, during their first 12 years of life, junior recruits from broods with either low or high levels of hatching asynchrony should breed less frequently, show lower breeding longevity and achieve lower accumulated breeding success.

## Methods

### Study population

Blue-footed boobies of Isla Isabel, Nayarit, Mexico (21°50’59.0"N 105°52’54.0"W) can live up to 25 years (Ortega et al., 2017) and start reproduction between their first and twelfth years of life (Drummond et al., 2011). Females lay 1-3 eggs per nest –with a median clutch of two (Ortega et al. unpublished data)–, and parents begin incubation immediately after the laying of the first egg. Two-egg clutches are the most common clutch size followed by one-egg clutches, representing 59% and 30%, respectively, of 33,975 nests established between 1989 and 2022. During these 33 years of monitoring, two- and one-chick represented 44% and 47%, corresponding,ly of 17,093 broods. Two-chicks broods often fledge both nestmates (∼61%, while 22% of them only fledge one chick, Ortega et al., unpublished data). Nests with more than three eggs or nestlings are rare, representing fewer than ∼0.00% of all nests.

### Demographic data

From 1989 to 2022, breeders within two study areas of the Isla Isabel colony were individually identified by alphanumeric steel bands fitted when they fledged (reached age 70 days). Throughout each field season (roughly February through July), all pairs were monitored by recording their nest contents every 3 days through fledging (details in Drummond et al., 2003). Depending on their hatching order, nestlings were temporarily identified with plastic bands. This monitoring protocol allowed us to estimate both laying and hatching dates within a two-day error margin by assuming that laying/hatching occurred two days prior to first sighting of an egg/hatchling (Drummond and Rodríguez, unpublished data). To minimize disturbance to incubating birds, eggs were not marked and therefore laying order cannot be determined. Laying dates of nestlings present at the start of monitoring were estimated by subtracting their age –estimated from beak and ulna growth curves– from the date when monitoring began (Drummond and Rodríguez, unpublished data).

Daily monitoring in 1985 and 1990 of a sub-set of clutches laid before and during monitoring allowed us to tally both laying and hatching asynchrony of 70 two-egg clutches more precisely. Median laying asynchrony was of 5 days with a median absolute deviation (MAD) of 1 day and range of 1-14 days, while median hatching asynchrony was of 4 days with a MAD of 1 day and range of 1 0-13 days, (Figure 2a). Furthermore, the median laying asynchrony of 672 nest (all two-egg clutches and two-chicks broods) established during the standard monitoring –and thus did not require estimations of laying dates– between 1989 and 2022 was of 3 days with a MAD of 0 days and ranged from 0 to 18 days.

**Figure 1.**
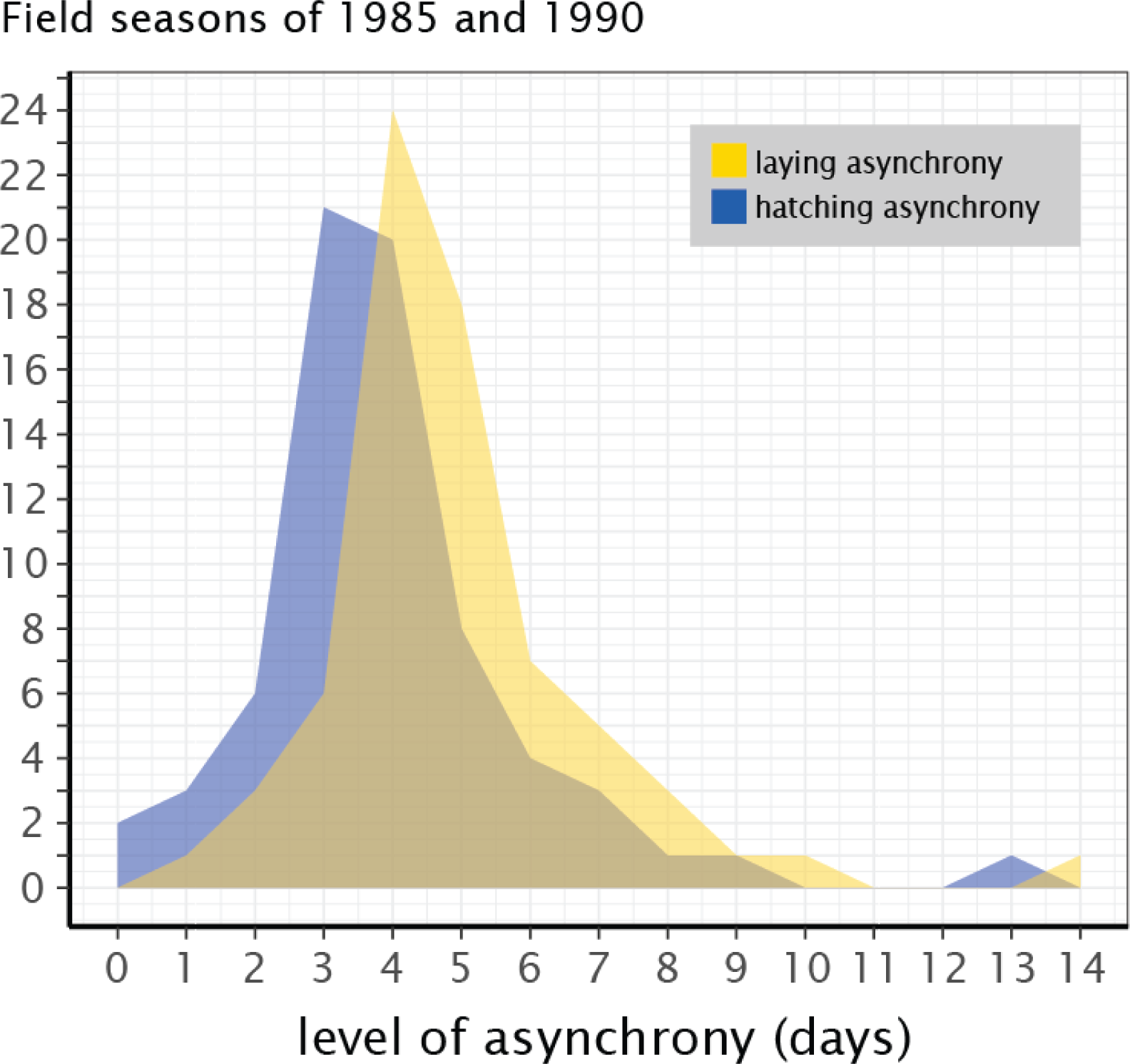
Frequency of laying and hatching asynchronies of 70 daily monitored two-egg clutches in which both eggs hatched.

**Figure 2.**
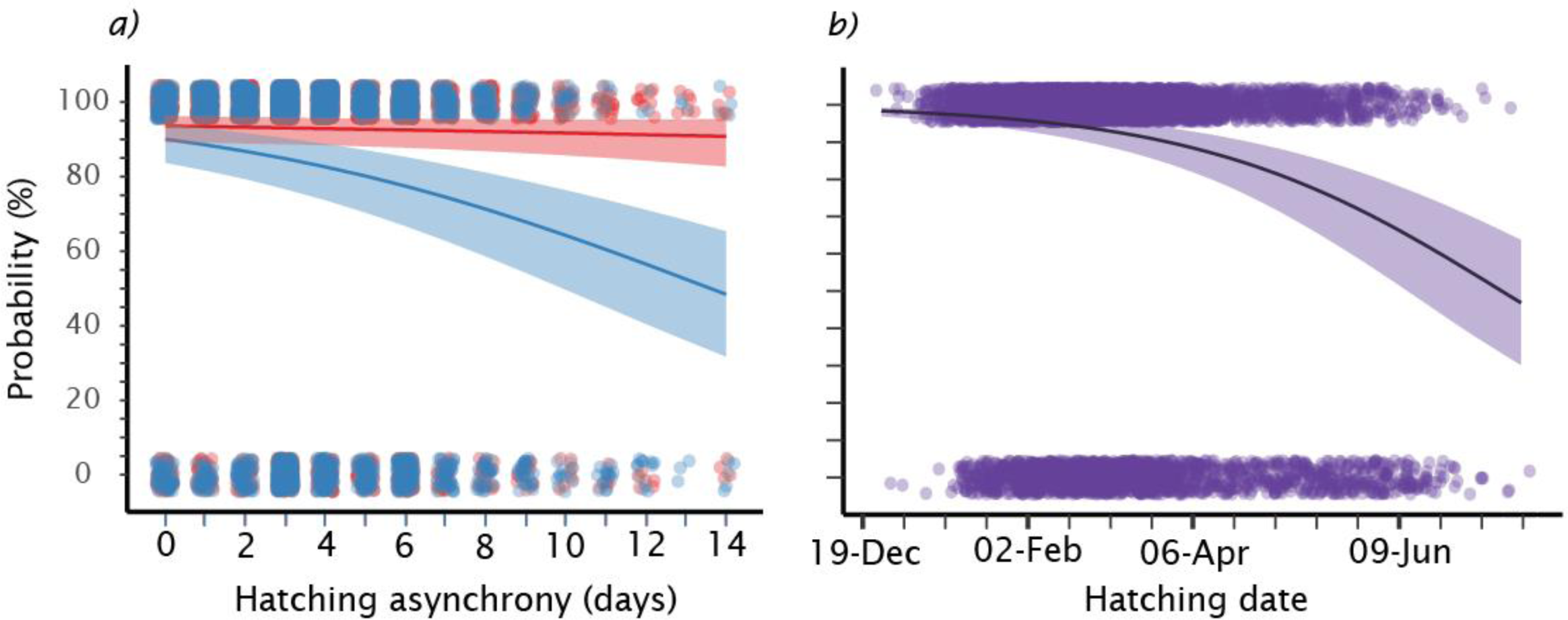
Fledging probability of 6645 nestlings from 3331 two-chick broods in response to a) the interactive effect of hatching asynchrony and hatching order and b) hatching date. Senior and junior offspring are shown in red and blue, respectively. Median effects and their 89% highest posterior density intervals are presented as shaded areas; jittered dots are raw observations.

**Figure 3.**
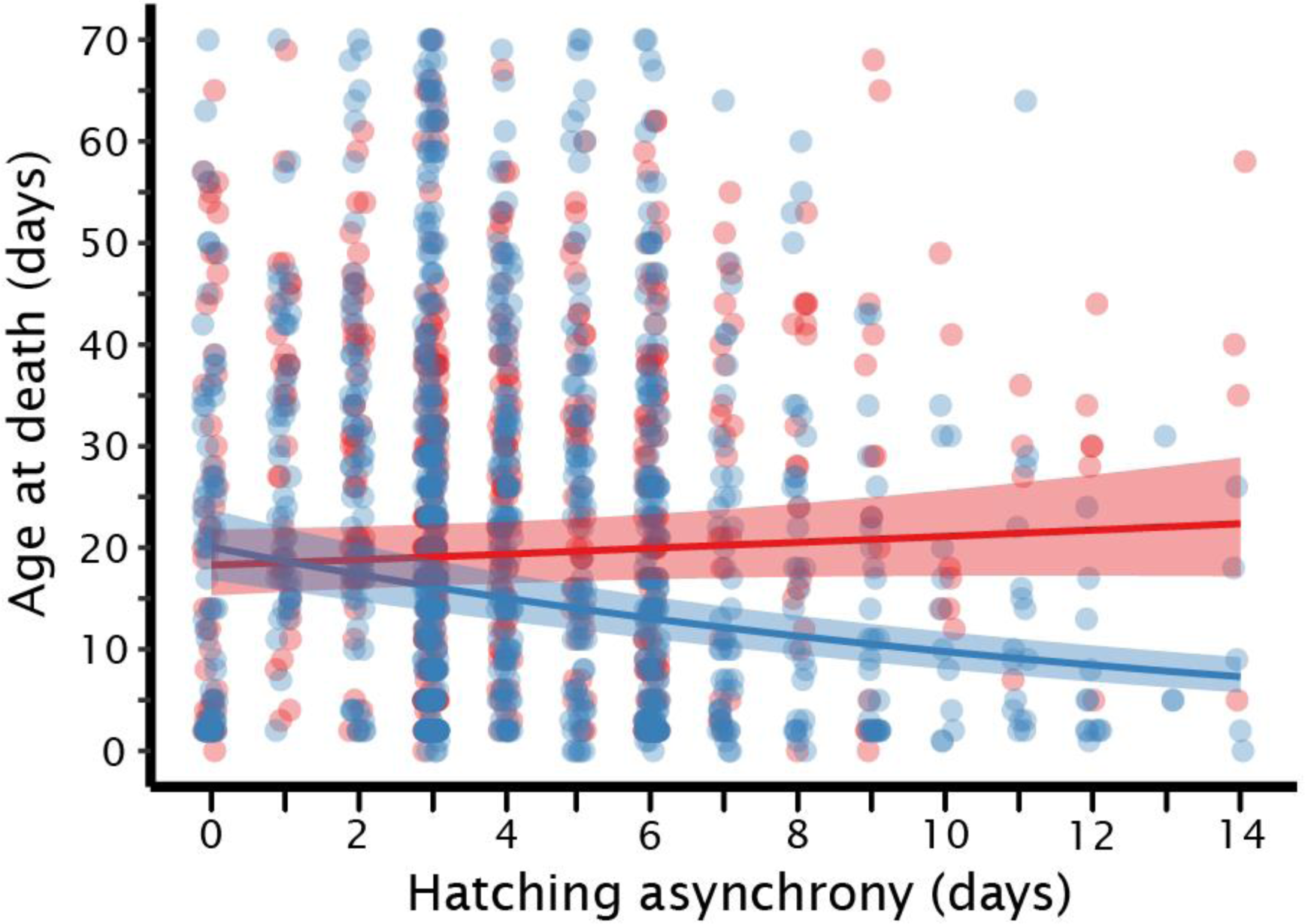
Synergetic effect of hatching asynchrony and hatching order on age at death of 641 senior and 966 junior nestlings that died in 1134 broods of two. Senior and junior nestlings are displayed in red and blue, respectively. Median effects and their 89% highest posterior density intervals are presented as shaded areas; jittered dots are raw observations.

### Fitness consequences of hatching asynchrony

Using all clutches where two chicks hatched between 1990 and 2022, we assessed the fitness consequences during the nestling period of hatching asynchrony (across its naturally occurring span from 0 to 14 days). Particularly, we tested for effects of hatching asynchrony (absolute difference between estimated hatching dates of senior and junior siblings) on a) probability of fledging, (whether nestlings reached 70 days old, b) age at death of nestlings that died before 70 days, and c) body weight at fledging. Our sample excluded clutches with >14 days of hatching asynchrony, as these high values might have arisen from the differences in hatching dates between first- and third-hatched offspring of nests were the second-laid egg was lost and was not recorded (Drummond, personal observations).

Using all fledglings from two-chick broods born between 1990 and 2010, we tested for consequences of hatching asynchrony for five late-life fitness traits: a) recruitment probability, that is, breeding in the study area within 12 years (later recruitment is unlikely; Drummond et al., 2011), b) age at first reproduction, c) number of breeding events, d) accumulated breeding success, and e) age at last sighting (a proxy of reproductive longevity). Accumulated breeding success was the sum of all fledglings produced. Median hatching asynchrony for the 6645 nestlings belonging to 32 birth cohorts was 3 days with a MAD of 1 day. Median body weight, ulna length and beak length of the fledglings were 1600 g with a MAD of 180 g, 204 mm with a MAD of 9 mm, and 106 mm with a MAD of 4 mm, respectively.

### Ethical note

Permission for fieldwork was given by Mexican authorities (SEMAR, CONANP, SEMARNAT and Parque Nacional Isla Isabel), following the ASAB /ABS (2023) Guidelines for the treatment of animals. During our annual monitoring, two volunteers visually inspected nests and parents (without handling them). To reveal nest contents and to read the incubating bird’s steel ring, one volunteer carefully nudged the adult backward with a forked stick. After withdrawing from the nest, the volunteers waited nearby until eggs and young chicks were covered by a parent and, if the parent flew from its nest, covered the eggs with leaves to prevent predation from overflying gulls.

#### Statistical analyses

All analyses were performed in the R statistical environment (R Development Core Team, 2022). All continuous fixed variables were standardized prior to model fitting to facilitate the understanding of parameter estimates (Cade, 2015; Grueber et al., 2011). Weakly informative priors were included in the analyses to provide regularization to stabilize computation and to avoid overfitting and erroneous estimations of large effect sizes (Gelman et al., 2008; Lemoine et al., 2016). A normal prior of N(0,1) was allocated to the fixed effects, which implies that we expect most responses to be within one standard deviation of the median response value and that large effects should be unusual (Lemoine et al., 2016). The posterior distribution of the parameters, alongside their 89% highest posterior density intervals (HPD) –a high-probability interval of parameter values (McElreath, 2020)–, were drawn by running four randomly initiated Markov chains. Each Markov chain ran for 10,000 iterations with a burn-in of 1000 iterations. Posterior predictive checks were carried for each generalized linear mixed models (GLMMs) using the function *launch_shinystan* of the package shinystan (Gabry, Veen, et al., 2022). The *rescale* function of the package *arm* (Gelman et al., 2016) was used for variable standardization. All GLMMs were built using the *stan_glmer* function in the *stanarm* package (Gabry, Ali, et al., 2022).

For all models, natal nest ID along with the focal bird’s mother’s ID were added as random effects, since siblings, which are statistically non-independent, were included in the sample. Furthermore, birth cohort was added as random effect to account for any unmeasured environmental conditions (e.g., snake predation, the interactive effects between oceanic and atmospheric phenomena; (Ortega et al., 2021). Pertinently, the linear expression of hatching asynchrony tested whether greater asynchrony benefits the senior sibling while negatively affecting its sibling, for example, by rapidly disposing of their sibling, senior offspring could dodge any direct fitness penalties derived from sharing parental investment. The quadratic expression of hatching asynchrony also tested for the possibility that the increased aggression between nestmates in more synchronous broods could also hinder the fitness of either nestmate. As nestlings/juveniles are not sexed until they recruit, the analyses of early-life survival, weight at fledging and recruitment probability did not include sex as a covariate.

### Fitness consequences in the nestling period

To analyse fledging probability, age at death, and fledgling weight we fitted a binomial (logit link function), negative binomial (log link function), and Gaussian (identity link function) models, respectively. The sample for the fledging probability analysis comprised 6645 nestlings (3321 and 3324 seniors and juniors, respectively) from broods of two. For age at death, we used data from 641 senior and 966 junior nestlings that died before fledging (≤ 70 days). Measures of body weight, ulna and beak length were available for 4590 nestlings (2456 and 2134 seniors and juniors, respectively) and these comprise the sample for the fledgling weight analysis. All models shared the same base structure, containing hatching order, the linear and quadratic expressions of maternal age, and each offspring’s hatching date as fixed effects. Alongside their base model, six competing models were fitted for each dependent variable, testing for both the linear and quadratic effects of hatching asynchrony, plus their corresponding two-way interactions with hatching order and with hatching date. The base model structure for the fledging weight analysis also included ulna and beak length as fixed effects.

Hatching order (a two-level categorical variable: first-hatched, and second-hatched), accounted for nestling mortality, which, in this population, increases with hatching order (Torres & Drummond, 1997). Hatching date (standardized by setting November 3^rd^ as day 1) controlled for intra-seasonal weather variations: in Isla Isabel, sea surface temperature increases and rainfall levels drop as the breeding season progresses, reducing primary ocean productivity, which results in higher mortality of chicks from nests established later in the season (Ortega et al., 2022). The quadratic expression of maternal age was added to account for reproductive senescence, which occurs in this species after age ∼10 years (Beamonte-Barrientos et al., 2010). Ulna and beak length controlled for skeletal size. The interactions between hatching asynchrony and hatching order were included to test whether greater levels of asynchrony affect last-hatched individuals more. The interactions between hatching asynchrony and hatching date tested for the possibility that poor natal weather conditions exacerbate the effects of hatching spans on last-hatched nestlings.

#### Fitness consequences in the juvenile and adult periods

To examine recruitment probability, age at first reproduction, number of breeding events, accumulated breeding success, and age at last sighting we built models with binomial (logit link function), negative binomial (log link function), and Poisson (log link function) error distributions. The sample for the recruitment probability comprised 1451 senior and 1258 junior juveniles whose first reproduction could be tallied within a period of 12 years. For analyses of age at first reproduction, number of breeding events, accumulated breeding success, and age at last sighting, we used the demographic data of 542 senior and 405 junior recruits. All analyses shared the same model structure and included hatching date and hatching order as fixed effects. For each dependent variable, six more competing models were fitted, testing for both the linear and quadratic effects of hatching asynchrony, plus their corresponding two-way interactions with hatching order and with hatching date. In this context, hatching order controlled for possible variations in fledging production between senior and junior individuals (e.g., Drummond & Rodríguez, 2013), while hatching date accounted for potential effects of adverse natal weather conditions (e.g., Ancona and Drummond 2013). Furthermore, two-way interactions between sex and either the linear or quadratic effects of hatching asynchrony were fitted for all dependent variables except recruitment probability. Age at first reproduction and age at last sighting were added as continuous fixed effects in the number of breeding events and accumulated breeding success. In the age at last sighting analyses, only age at first reproduction was added as a covariable. Age at first reproduction and age at last sighting were added to controlled for selective appearance and disappearance, respectively (van De Pol & Verhulst, 2006).

#### Model selection

To find each response variable’s best-supported model, we first estimated, for every model, the leave-one-out cross-validation information criterion (LOO-IC) to calculate the expected log predictive density (ELPD), which outlines the predictive fit of a model. The model with the highest ELPD value was selected (Hollenbach & Montgomery, 2020). But, if the difference in ELPDs between the best and next best supported models was not at least twice the estimated standard error of this difference, the models were considered equivalent (Hollenbach & Montgomery, 2020) and the most parsimonious one was selected. In the Supplementary Information Material, we provide the model selection tables containing all competing models’ LOO-IC and ELPD differences along with their estimated standard errors (Table S1-S3). LOO-IC and ELPD were estimated using the *loo* package (Vehtari et al., 2017).

## Results

### Fitness consequences in the nestling period

In two-chick broods, fledging probability was better explained by the model containing the two-way interaction between the linear expression of hatching asynchrony and hatching order (Table S1a). The fledging probability of first-hatched individuals was consistently high (∼93%), while that of second-hatched birds decreased from 90% to ∼48% between 0 and 14 days of asynchrony (Table 1; Fig. 2a). Fledging probability was affected by maternal age quadratically, progressively increasing from 70% at 1 year to 94% at age 9 years, plateauing between 9 and 12 years, and declining to ∼38% at 22 years (Table 1).

**Table 1.**
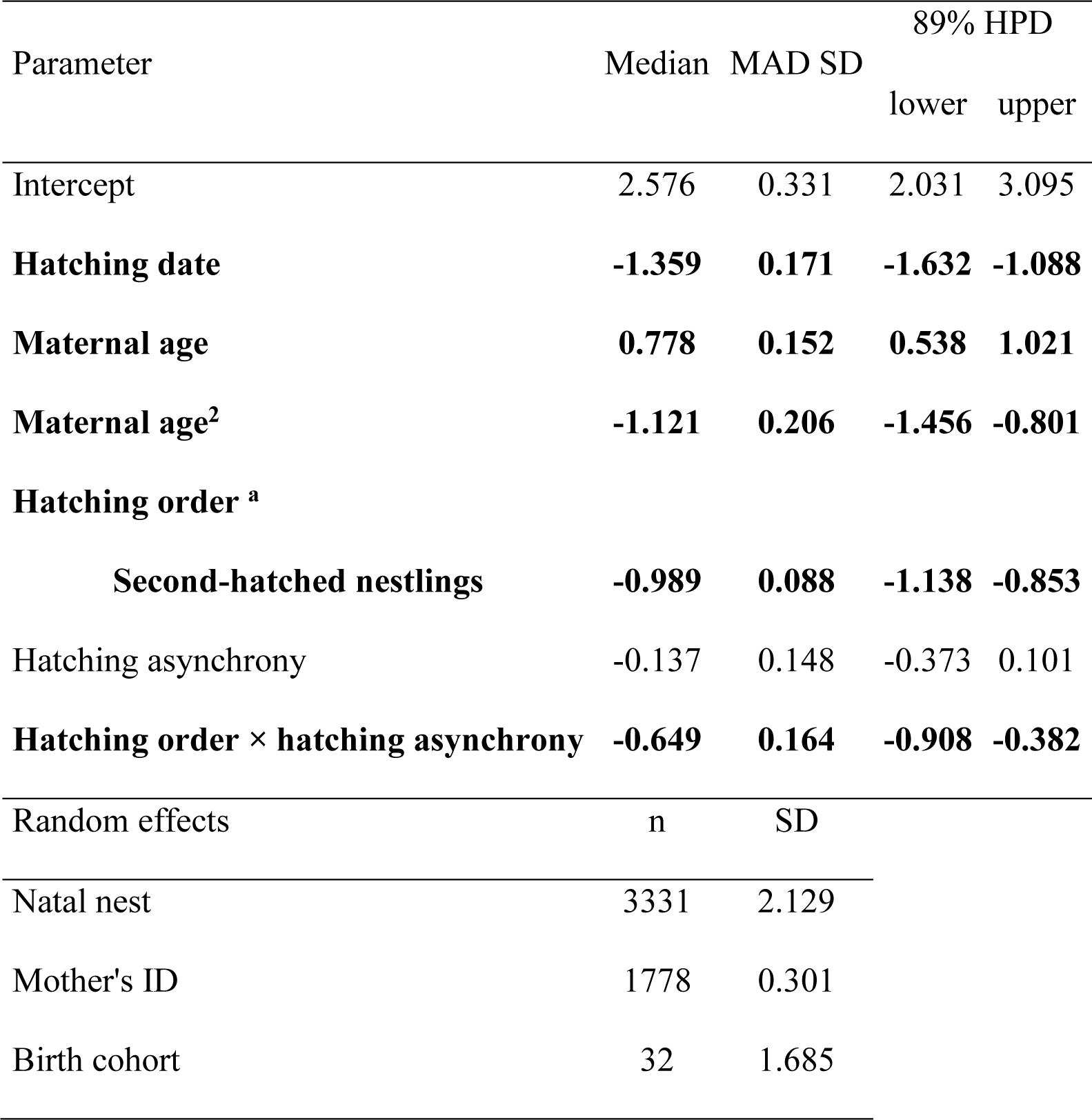
Effect of hatching asynchrony on the fledging probability of 6645 nestlings.

Furthermore, nestlings that hatched in mid-December (boreal Winter) had a higher median fledging probability than their counterparts hatched in early-July, that is, in boreal Summer (∼98% vs 47%; Table 1; Figure 2b).

Parameters whose highest posterior density (HPD) intervals did not contain zero are shown in boldface type. Median absolute deviations (MAD) from the standard deviation are provided (SD). ^a^ First-hatched nestlings were used as reference levels.

The age at which nestlings in broods of two died was explained by a two-way interaction between hatching asynchrony and hatching order (Table S1b). At zero days of asynchrony, both senior and junior nestlings died at similar ages (∼19 days old), but as asynchrony grew age at death increased for the former and decreased for the latter (∼ 22.3 and ∼ 7.3 days old at 14 days of asynchrony, respectively; Table 2; Fig 4). Maternal age affected nestlings’ age at death (Table 2), following an inverted-U pattern, increasing from ∼15.4 days at maternal age of 1 year to ∼19.7 days when the mother was 9 years old, then declining to ∼10.5 days at maternal age of 21 years. Critically, because avian fledging weights often predict fitness, weight of the (unsexed) nestlings at fledging was unrelated to either level of hatching asynchrony or hatching date (Table S1c). Regardless of maternal age or hatching date, both senior and junior fledglings had an estimated median weight of ∼1565 g (Table 3).

**Figure 4.**
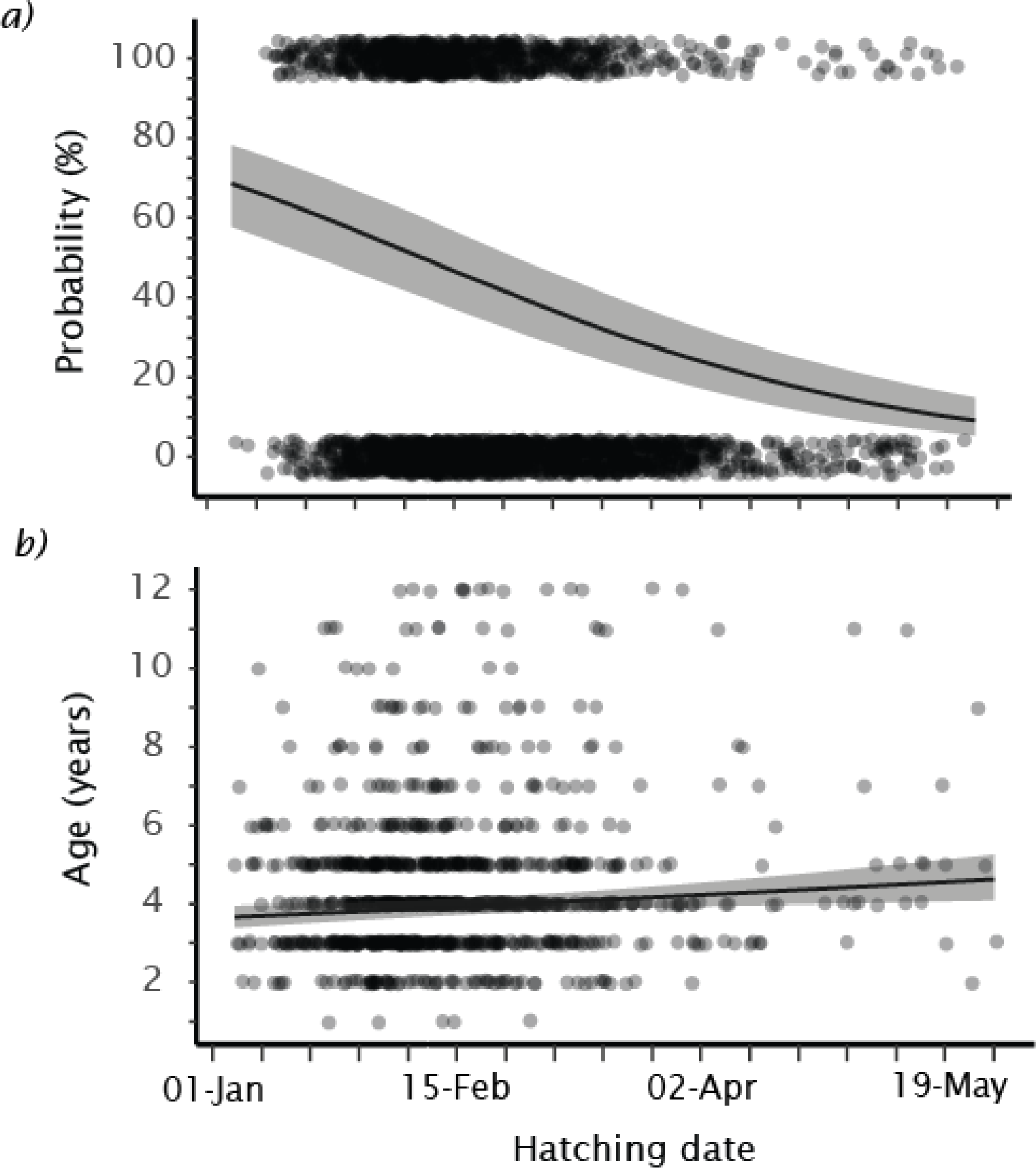
Effect of hatching date on a) recruitment probability of 1451 senior and 1258 junior juveniles and b) age at first reproduction of 542 senior and 405 junior recruits. Median effects and their 89% highest posterior density intervals are presented as shaded areas; jittered dots are raw observations.

**Table 2.**
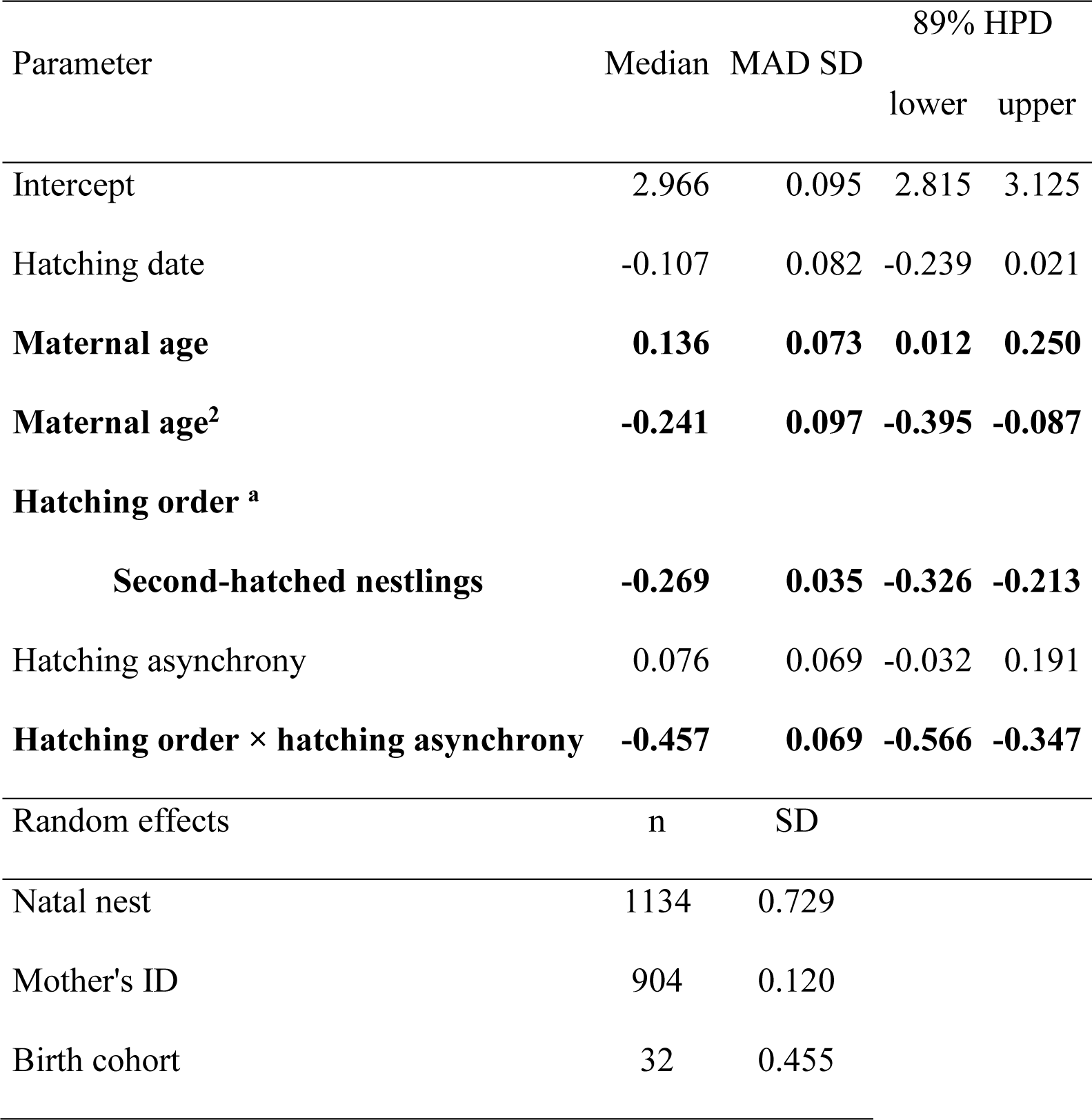
Effect of hatching asynchrony on age at death of 1607 nestlings in 1134 two-chick broods.

**Table 3.**
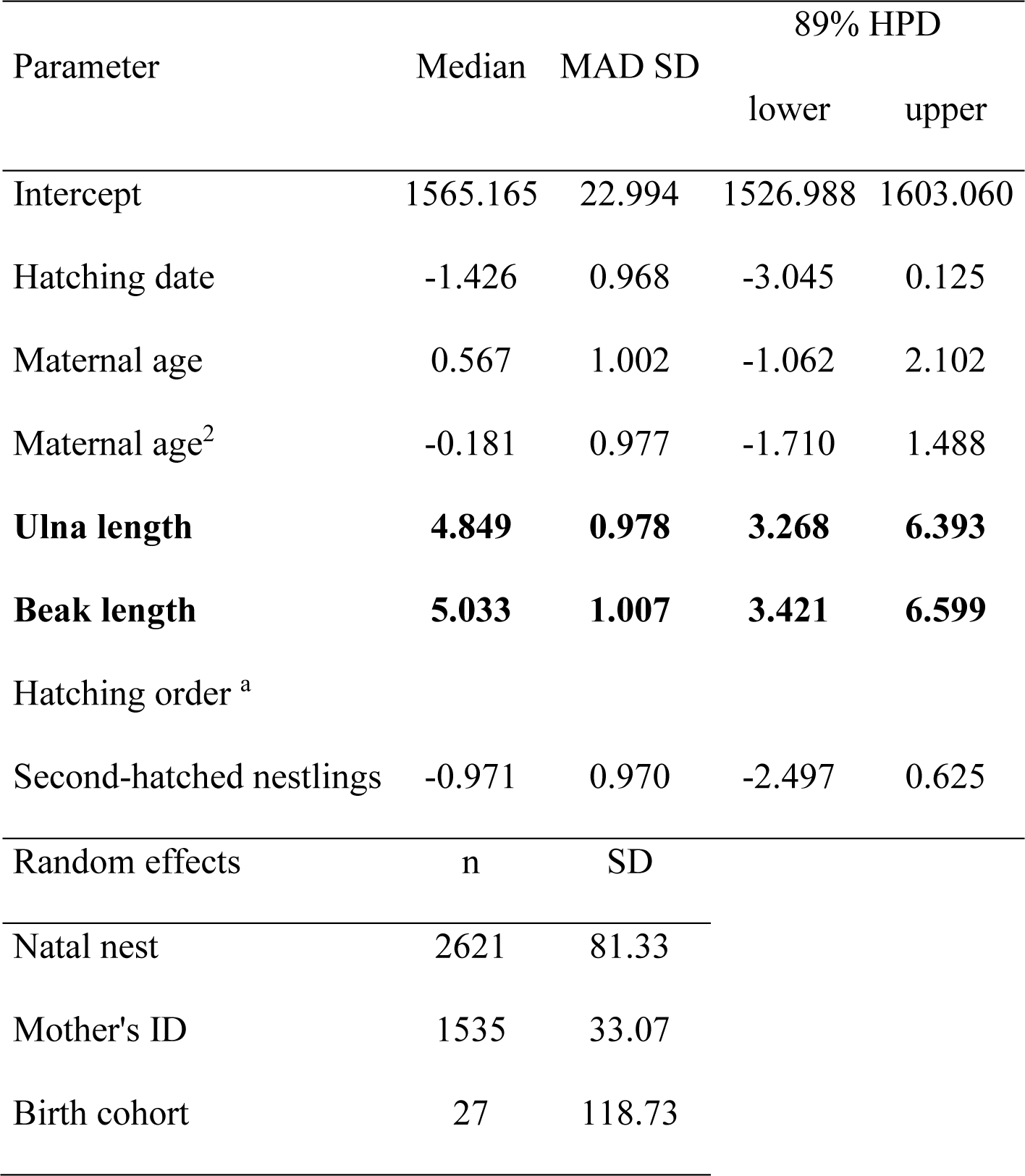
Effects of hatching asynchrony on the wsseight of 4590 fledglings from 2621 two- chick broods.

Terms whose highest posterior density (HPD) intervals did not contained zero are shown in boldface type. *MAD* Median absolute deviations. ^a^ First-hatched birds were used as reference levels.

Fixed effects whose highest posterior density (HPD) intervals did not included zero are shown in boldface type. *MAD* Median absolute deviations. ^a^ First-hatched nestlings were used as reference levels.

#### Fitness consequences in the juvenile and adult periods

No long-term fitness consequences of level of natural asynchrony were detected. In the sample of 1451 senior and 1258 junior juveniles, level of hatching asynchrony did not correlate with recruitment probability (Table S2). Nor for 542 senior and 405 junior recruits, did level of hatching asynchrony correlate with age at first reproduction (Table S3a), number of breeding events (Table S3b), accumulated breeding success (Table S3c), or age at last sighting (Table S3d).

However, both hatch order and hatch date had effects in the juvenile and adult periods. Median recruitment probability of first-hatched juveniles was ∼6% higher than of second-hatched juveniles (41.7% vs 35.6%, correspondingly; Odd ratio = 1.29 [89% HPD: 1.10 to 1.49); and for 405 junior recruits, median age at first reproduction was ∼0.25 years later than for 542 seniors (4.54 years vs 4.29 years, respectively; Odd ratio = 1.06 [895 HPD: 1.01 to 1.11]).

Early hatched juveniles were considerably more likely to recruit than their later-hatched counterparts (early-January and late-May hatched birds had 69% and 0.09% chances of recruiting, respectively; Table S4a; Figure 4a). Moreover, recruits that hatched late in the season recruited ∼0.97 years later than their early-hatched counterparts (Table S4b; Figure 4b). Finally, males recruited ∼0.81 years later than females (Odd ratio = 1.20 [89% HPD: 1.14 to 1.26]; Table 4Sb), at median ages of 4.84 and 4.03 years, respectively.

However, despite senior juveniles being more likely to recruit than juniors and recruiting earlier than juniors, senior and junior recruits did not differ in their number of breeding events (∼ median of 3.58 events), nor in their age at last sighting (median of ∼ 8.41 years old), regardless of their sex and hatching date. However, senior recruits (n = 542) produced 0.19 more fledglings in their first 12 years of life than junior recruits (n = 405) (2.38 vs 2.19 fledglings; Odd ratio = 1.09 [89%HPD: 1.02 to 1.16]: Table S4c).

## Discussion

We found effects of both hatching order and hatching asynchrony level on the early survival of senior and junior nestlings. Specifically, the greater the hatching asynchrony, the less likely were juniors to fledge. Moreover, while deaths of seniors and juniors in synchronous broods occurred at similar ages, they diverged increasingly as hatching span increased, juniors dying ever younger than seniors. Surprisingly though, seniors and juniors that fledged did not differ in body weight, independent of hatching asynchrony level.

Although we found long-term effects of hatching order, we found no long-term effects of hatching asynchrony level. Junior fledglings recruited slightly less often than seniors, their recruitment came 0.25 years later, and they fledged 0.19 fewer offspring in their first 12 years of life. Given the developmental susceptibility of birds to experimental stresses in infancy, including food scarcity and poor growth, compensatory growth, elevated corticosterone and parasitic infection, as well as poor weather, habitats, prey availability, parental care and late fledging (Drummond & Ancona, 2015), it is remarkable that there was no evidence of long-term fitness penalties of hatching second are any greater at the lowest and highest levels of asynchrony (our prediction) or that they increase linearly with level of hatching asynchrony. As far as we can tell, level of hatching asynchrony affects booby offspring only during the nestling period.

### Fitness consequences in the nestling period

As expected, a higher level of hatching asynchrony (above 0 days) was correlated with lower fledging probability and increased mortality at younger ages in junior birds. Increased agonistic interaction between siblings and disparity in motor skills may explain these patterns. In this booby, experimentally enlarged age/size disparities result in an 5-fold increase in aggression from the bigger nestling towards its smaller nestmate during the first 10 days of interaction (Osorno & Drummond, 1995). Similarly, in the black-legged kittiwake *Risa tridactyla*, junior chicks from broods with experimentally increased hatching asynchrony were attacked more often by their seniors during the first 10 days of life and also experienced a higher mortality throughout the nestling period than their seniors (Merkling et al., 2014). Thus, larger hatching spans, which can exacerbate aggression by senior offspring, can be detrimental to the junior offspring early survival.

Hatching synchrony appears to benefit juniors, as they can reach a fledging probability and weight like that of their older siblings (this study). An increase in aggressive sibling competition through synchronous hatching appears to act as a stimulus that elicits an increase in parental effort. For example, in experimental pairs of similar sized blue-footed offspring –which compete more aggressively– have been shown to receive 25% more food from parents during their first 20 days of life (Osorno & Drummond, 1995)). Similarly, parental feeding frequency has been showed to increase after 10 days of intense sibling aggression in experimentally synchronous broods of the black-legged kittiwake (Merkling et al., 2014). It remains to be seen if increasing levels of hatching asynchrony can alleviate the parental trade-off between current and future reproduction by offsetting the offspring’s food demand curves (Hussell, 1972),

#### Fitness consequences in the juvenile and adult periods

We did not detect any fitness consequences of the level of hatching asynchrony in the juvenile or adult periods of the blue-footed booby. This apparent lack of long-term fitness penalties can be attributed to the filtering out of junior nestlings from more asynchronous broods and juveniles that hatched later in the season. Additionally, developmental resilience, which probably relies on increased investment by parents, physiological adaptations, and life history adjustments by the offspring, might explain why some of the junior nestmates from asynchronous nests and individuals that hatched later in the season (i.e., those that faced adverse natal weather conditions) survived and showed no fitness deficits later in life.

In the blue-footed booby, advancement of recruitment age by fledglings raised in a year under ENSO-like conditions –lack of rainfall and warm waters (Magaña et al., 2003)– is thought to contribute to the apparent lack of silver spoon effects, as it seems to allow these individuals to maximize their long-term fitness (Ancona & Drummond, 2013). Interestingly, junior juveniles did not recruit earlier than senior juveniles (or *vice versa*). However, both juniors and juveniles that hatched later in the season delayed their recruitment, as expected for a species with a slow pace of life (i.e., those with low reproductive rates, slow-developing offspring, and long lifespans; (Gaillard et al., 1989)), which tend to postpone breeding until they are able to face the costs of reproduction (Weimerskirch, 1992). Thus, by delaying their first reproduction, junior and offspring that hatched later in the season can match both seniors and birds that hatched earlier in the season in terms of number of breeding events and age of last sighting. Physiological trade-offs in early-life (e.g., (Stier et al., 2015)) could also play a key role in diminishing the fitness impacts of greater levels of asynchrony and adverse natal weather condition on second-hatched individuals and later-hatched offspring, respectively.

Regardless of the level of hatching asynchrony, junior recruits produced 0.19 fewer fledglings than senior recruits during their first 12 years of life, a minor difference indicating suboptimal condition in early adult life. Indeed, second-hatched female boobies are ∼8% underweight compared to first-hatched females during their first reproductive events at ages 4-6 years (Carmona-Isunza et al., 2013). Moreover, junior recruits of both sexes start reproducing later in life than senior recruits (this study). Despite their overall developmental resilience, the post-hatching social environment appears to take a toll on junior recruits. Nonetheless the importance of junior recruits’ 0.19 deficit in fledgling production of second-hatched boobies is of biologically relevant remains to be seen. Tallying of eventual recruitment of the fledglings produced will be required to plumb the fitness consequences of both hatching asynchrony and hatching order.

Here we showed for a population of blue-footed boobies off the Eastern-Pacific Coast that the level of hatching asynchrony affects only early-life survival of junior nestlings. From these offspring point of view, hatching synchrony is beneficial, as it appears to allow them to dodge early-life fitness penalties, presumably though increased parental investment during the rearing stage. Our results provide some insight into how parentally imposed natal social environments affects short- and long-term fitness in wild populations. Further studies should focus on how the level of hatching asynchrony impacts the parents’ fitness, as trade-offs are expected to occur when increasing current investment, particularly so for species with a slower pace of life. The factors that elicit both laying and hatching spans should also be explored, particularly if they affect parental survival.

## Author Contributions

**Santiago Ortega**: Conceptualization, Investigation, Original draft, Methodology, Formal analysis, Data presentation. **Cristina Rodríguez**: Investigation, Revision of manuscript, Data curation. **Hugh Drummond**: Investigation, Resources of funding, Revision of manuscript, Administration, Supervisions of project.

## Data Availability

All data analysed in this study are included as Supplementary material.

## Declaration of Interests

We declare no conflict of interest.

## Supporting information

supplementary material database

## Acknowledgements

We thank the Armada de México and the Comisión Nacional de Áreas Protegidas for their logistical support of annual fieldwork. We are thankful for the many volunteers and the fisherman of Isla Isabel for assistance in the field. S.O thanks and acknowledge the financial support of the Consejo Nacional de Ciencia y Tecnología during the study.

## Notes

### Competing Interest Statement

The authors have declared no competing interest.

